# Noise-induced bistability in the quasineutral coexistence of viral RNA under different replication modes

**DOI:** 10.1101/272906

**Authors:** Josep Sardanyés, Andreu Arderiu, Santiago F. Elena, Tomás Alarcón

## Abstract

Evolutionary and dynamical investigations on real viral populations indicate that RNA replication can range between two extremes given by so-called stamping machine replication (SMR) and geometric replication (GR). The impact of asymmetries in replication for single-stranded, (+) sense RNA viruses has been up to now studied with deterministic models. However, viral replication should be better described by including stochasticity, since the cell infection process is typically initiated with a very small number of RNA macromolecules, and thus largely influenced by intrinsic noise. Under appropriate conditions, deterministic theoretical descriptions of viral RNA replication predict a quasineutral coexistence scenario, with a line of fixed points involving different strands’ equilibrium ratios depending on the initial conditions. Recent research on the quasineutral coexistence in two competing populations reveals that stochastic fluctuations fundamentally alters the mean-field scenario, and one of the two species outcompetes the other one. In this manuscript we study this phenomenon for RNA viral replication modes by means of stochastic simulations and a diffusion approximation. Our results reveal that noise has a strong impact on the amplification of viral RNA, also causing the emergence of noise-induced bistability. We provide analytical criteria for the dominance of (+) sense strands depending on the initial populations on the line of equilibria, which are in agreement with direct stochastic simulation results. The biological implications of this noise-driven mechanism are discussed within the framework of the evolutionary dynamics of RNA viruses with different modes of replication.

## I. INTRODUCTION

An essential yet poorly understood process during intracellular amplification of viral genomes is the mode of genome replication. Very few theoretical studies have explored the dynamical properties of alternative modes of RNA virus genomes replication [1–3], and even fewer experimental studies have collected data properly describing the temporal increase of viral sense genomes and of antisense genomes that are the unavoidable intermediates of replication [4]. Nevertheless, the scarce available data suggest different models of replication for different viruses. Two extreme mechanisms of RNA virus genome replication have been discussed. The first mechanism is referred as the stamping machine replication (hereafter SMR) mode. For SMR, and considering an infecting virus of (+) sense RNA genome, the whole progeny of (+) sense strands will be synthesized from a few (−) sense strands complementary to the infecting (+) one. In SMR, an asymmetric accumulation of strains of both polarities exists: (+) strands are massively produced while intermediate (−) strands stay at low concentration. Under this mode of replication, the expected fraction of mutant genomes produced per infected cell follows 1 − *e*^−*μ*^, being *μ* the genomic mutation rate. In this case, the distribution of mutants per infected cell before the action of selection follows a Poisson distribution. Such a distribution of mutants has been described for bacteriophages *ϕ*X174 [5] and Q*β* [6]. The second possible mode of replication is the geometric replication (hereafter GR). For GR replication is symmetrical and both (+) and (−) sense RNA strands are used as templates for viral amplification with equal efficiency. For this mode of replication, the expected fraction of mutants genomes produced per infected cell depends on the number of replication cycles, *n*, according to expression 1 − *e*^−*nμ*^. The resulting distribution of mutants then follows the Luria-Delbriick distribution. Deviations from the Poisson distribution were found for the phage *T*2 [7], thus suggesting that such a virus replicates according to the GR model. In principle, these two modes of replication represent the two extremes of a continuous of possible strategies. In this sense, intermediate modes of replication have been described for bacteriophage *ϕ*6 [8] and turnip mosaic potyvirus (TuMV) [4]. In these cases, the distribution of mutants slightly deviated from the Poisson distribution, thus suggesting that the replication was mainly achieved by an SMR strategy plus a small contribution of GR.

Results from experiments specifically designed to determine the mode of replication for eukaryotic RNA viruses have been published only in recent years. Of particular interest, Martínez et al. [4] monitored and quantified the accumulation of both RNA polarities of TuMV infecting *in vitro* protoplasts of the host plant *Nicotiana benthamiana*. In this study, a simple dynamical model describing the production of (−) and (+) sense RNAs and interference of (+) strands on the synthesis of (−) ones was proposed (see Section II). The model was then used to fit the experimental data and to estimate replication parameters, most remarkably the fraction of molecules produced by SMR and GR. Inspired by this study, later on Combé et al. [11], working with the (−) sense RNA *Vesicular stomatitis virus* (VSV), and Schulte et al. [9], working with the (+) sense poliovirus, have shown that GR is the main mode of replication for these very different viruses.

Stochasticity is an essential component of molecular biosystems and so does virus replication. Firstly, infection is generally initiated by one or very few genomes per cell (a variable known as multiplicity of infection or MOI) which has to bind to cellular components to begin replication. Secondly, once starting to be synthetized, viral product have to find and interact with their appropriate cellular targets in a crowed environment. Accordingly, the outcome of an infection is likely to be affected by variability in the initial molecular interactions between virus and host cells and result into heterogeneity among infected cells (e.g., [10], [11]). Viruses have evolved mechanisms to minimize the effect of noise, such as for example actively controlling MOI [12] or limiting replication to well-defined and space-limited membrane-associated compartments known as viral replication factories [13].

The impact of stochasticity in the fate of replicator systems is a current subject of interest. Especially, the effects of noise on so-called quasineutral or degenerate scenarios, found in the deterministic limit. These scenarios involve a particular phase space topology in which quasineutral invariant sets appear (also called normally hyperbolic invariant manifolds [16]). Typically, these invariant sets display hyperbolicity in the normal direction, and the equilibrium state depends on the initial conditions [14, 15]. It is known that when noise is considered, the dynamics is dominated by the stochastic drift on this invariant manifold, and outcompetition of species due to stochasticity can be achieved [14, 15]. In the present article we analyze a simplified version of the model introduced in Martínez et al. [4], used to quantify the within-cell replication dynamics of (+) sense RNA viruses. This model displays a quasineutral coexistence scenario between (+) and (−) sense RNA strands when only considering strands’ replication and competition (see [3] for details). We analyze in detail the quasineutral dynamics of this model, focusing on the impact of stochastic drift on the invariant manifold which determines the fate of viral RNAs. We provide analytical approximations for the probabilities of achieving a given asymptotic state as a function of the initial conditions on the manifold. The analytical results are complemented with stochastic simulations. Finally, we describe a novel mechanism for noise-induced bistability.

## II. MATHEMATICAL MODELS

### A. Deterministic model

A simple mathematical model considering asymmetric RNA replication modes has been recently introduced in Ref. [3] (see also [4]). This model is given by the next couple of differential equations:

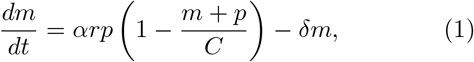

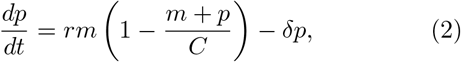

The system Eqs. (1)-(2) are deterministic rate equations of two strands of viral RNA with different polarities (*m*: minus and *p*: plus) that compete for the available resources and differ only in the time scales of their evolution (birth and death rates) defined by the densities amplification rates *αr* and *r*. Constant *C* is the carrying capacity. Since we are interested in the impact of the replication mode, we will focus our study on parameter *α*. When *α* = 1 replication proceeds via GR since both strands are replicated at the same rates. The cases *α* → 0 follow the SMR mode, since replication becomes asymmetric and there is much more production of (+) sense RNAs from the (−) one(s), which act as template(s). The two strands’ common carrying capacity *C* indicates the total number of individuals in the non-empty steady state.

Here we focus on the case *δ* = 0, which gives place to a particular topology of the phase space, given by a neutrally stable invariant line (see below). The deterministic continuum description is presumably applicable when *m* and *p* are 𝒪(*C*) and *C* >> 1. Let us, for convenience, introduce the scaled population variables *x* = *m*/*C* and *y* = *p*/*C* and rescale the time variables by the growth rate *r*, to write the system

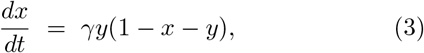

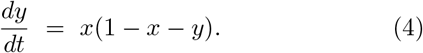

where the ratio of time scales is γ = (*αr*)/*r* = *α*.

### B. Stochastic model

A first level of description involves describing stochastic evolution of the integer-valued random processes *X*_*t*_ and *Y*_*t*_. Let us consider a Markov model given by the master equation (ME), which is obtained from the general form:

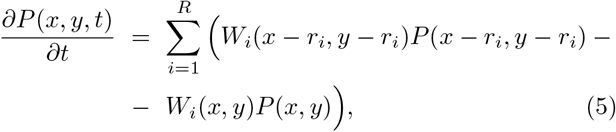

where *P*(*x*,*y*,*t*) is the probability of having *x* and *y* molecules at time *t*. *R* is the number of reactions, *W*_*i*_ and *r*_*i*_ are the transition rates (propensities) and the stoichiometry of the system. The ME for the system studied here, considering negative (*n*) and positive (*p*) sense strands can be obtained from the following reactions, after rescaling parameters:

1. Reactions for negative-sense strands:

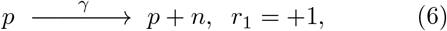

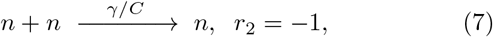

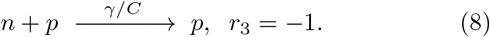
2. Reactions for positive-sense strands:

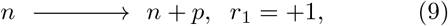

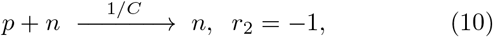

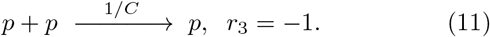

Reaction (6) denotes the synthesis of (−) sense strands from (+) sense ones. Reactions (7)-(8) involve the decrease of the population (*r*_2,3_ = −1), since they represent, respectively, competition between strands of the same polarity (intra-specific competition in ecological terms) and of different polarity (inter-specific competition). The same reactions are represented for (+) sense strands in reactions (9)-(11). Let us denote *n* and *p* as the number of (−) and (+) sense strands and *P*(*n*, *t*) and *P*(*p*, *t*) as the probability of having *n* and *p* strands at time *t*, respectively. The propensities for reactions (6)-(8) are given by *W*_1_ = γ*p*, *W*_2_ = (γ/*C*)*n*(*n* − 1), and *W*_3_ = (γ/*C*)*np*; while the propensities for reactions (9)-(11) are given by *W*_1_ = *n*, *W*_2_ = *C*^−1^*pn*, and *W*_3_ = *C*^−1^*p*(*p* − 1).

As mentioned, we consider a Markov model where the birth and death rates are, respectively, *β*_*y*_ and *β*_*x*_ and 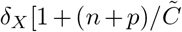 and 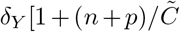 when *X*_*t*_ = *n* and *Y*_*t*_ = *p*, where 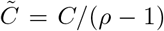 with *β*_*Y*_/*δ*_*X*_ = *β*_*X*_/*δ*_*Y*_ = *ρ* > 1.

This means that 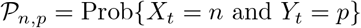 evolves according to the ME:

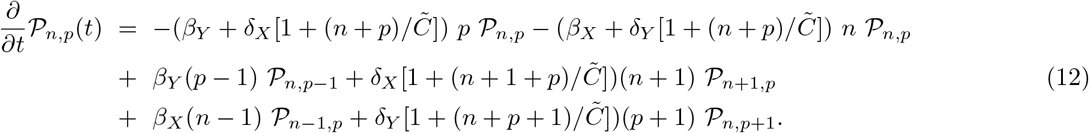

The low-density replication rates found in the deterministic differential equations are *αr* = *β*_*Y*_ − *δ*_*X*_ and *r* = *β*_*X*_ − *δ*\_*y*_, and the ratio of evolution time scales is γ = *δ*_*X*_/*δ*_*Y*_ = *β*_*Y*_/*β*_*X*_. Under suitable conditions, a diffusion approximation can be applied to the Master Equation (12), so that the variables 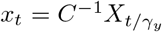 and 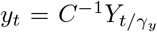 evolve according to the stochastic differential equations (SDEs):

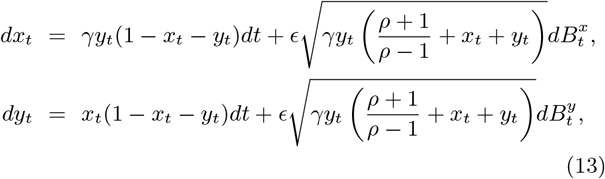

where 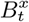 and 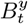 are independent Wiener processes and the noise amplitude is *ϵ* = *C*^−1/2^.

## III. RESULTS AND DISCUSSION

### A. Deterministic dynamics

As mentioned, we will focus our study on Eqs. (1)–(2) considering *δ* = 0. The parameters estimated in [4] from experimental data using Eqs. (1)-(2) revealed that the degradation rates of the viral RNAs were approximately between one and two orders of magnitude lower than the replication rate. Hence, the assumption *δ* = 0 is a first good approach to study the simplest model of viral strands amplification under differential replication modes. The fixed points and their stability for the system Eqs. (1)-(2) with *δ* = 0 was investigated in [3]. In the following lines we summarize these results from the scaled Eqs. (3)-(4). The fixed points for this system are given by the point 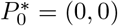 and the line of fixed points 𝕃* = (1 − *y*\*,*y*\*). The Jacobian matrix for this model is given by:

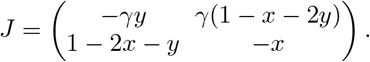

The local stability of 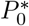 and 𝕃* can be studied from the eigenvalues obtained from the characteristic equation det 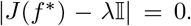, being *f*\* a fixed point and 𝕀 the identity matrix. The eigenvalues of the fixed point 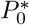 are 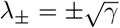. Thus the point (0, 0) is a saddle since γ > 0. The eigenvalues tied to the equilibria of the line 𝕃* can be computed by linearizing around (*x*\* = 1 − *y*\*, *y*\*). It can be shown that the eigenvalues obtained from det 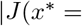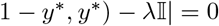 are given by

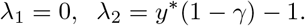

Note that for *y*\* ∈ [0,1] and γ ∈ (0,1] λ_2_ < 0, there exists a continuous line of marginally stable fixed points. Hence, the equilibria forming the line 𝕃* have a neutral eigenvalue (λ_1_, which is not locally attracting) and a negative eigenvalue (λ_2_, which is locally attracting). This gives place to a so-called normally hyperbolic invariant manifold [16]. These features involve that the initial conditions outside the line 𝕃* are attracted towards it and once they achieve the line they stop and remain there in forward time.

These dynamical characteristics can be visualized in Fig. 1. Figure 1b displays a phase portrait (obtained numerically using the 4th order Runge-Kutta method) for a case close to the SMR, using γ = 0.06. Note that different initial conditions end up in different places on the line 𝕃* place on the antiantidiagonal of the phase space [0,1] × [0,1]. The changes in the mode of replication are also important as shown in Fig. 1c. Here different initial conditions for (+) sense strands achieve equilibrium values on 𝕃* within the range 0.8 ≲ *p*(*t* → +∞) < 1 for the SMR (γ = 0.1), while for the GR (γ =1) the equilibrium values are found within the range 0.5 ≲ *p*(t → +∞) < 1.

**FIG. 1:**
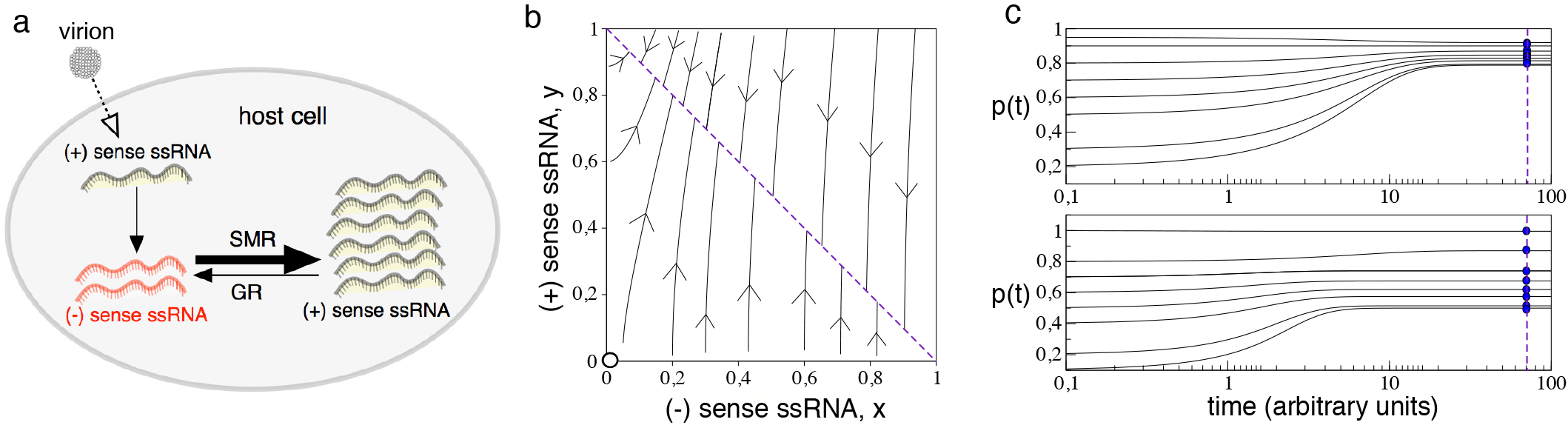
(a) Schematic diagram of asymmetric modes of RNA amplification for single-stranded (+) sense RNA viruses. Upon the infection by a virion, the entering (+) RNA is amplified following either the stamping-machine replication (SMR) or the geometric replication (GR) modes. The geometric replication involves that both (+) and (−) sense strands are produced at the same rates. Oppositely, for the SMR, a few (−) sense RNAs are used as the main template for an asymmetric replication, with a dominance of (+) sense RNAs. (b) Phase portrait displaying the quasineutral scenario obtained from the model proposed in [3] employed to study differential replication modes. Without degradation of RNA the equilibria states are a continuum of fixed points (dashed line in the antiantidiagonal), and the final populations depend on the initial conditions. Here we display this degenerate case using γ = 0.06, which is closer to the SMR model. (c) Time series (in log-linear scale) for (+) sense strands, *p*(*t*), starting from different initial conditions with γ = 0.1 (upper panel) and γ = 1 (GR, lower panel). The vertical dashed lines indicates the asymptotic states achieved by different initial conditions.

### B. Stochastic dynamics

In the previous section we have shown that 𝕃* is a neutral attracting line. It is known that this neutrality i.e., orbits achieving an invariant quasineutral line, is broken under the presence of noise [14, 15]. This means that, under stochasticity, the orbits are fastly attracted towards the line 𝕃*, but then stochastic fluctuations make the orbits to move along 𝕃*, ultimately achieving one of the two opposite states, given by either the point (1, 0) or (0,1) in the system studied here.

#### 1. Stochastic simulations and asymptotic dynamics

The dynamics of the master Eq. (12) is simulated using the Gillespie algorithm [17, 18] applied to reactions (6)-(11). Specifically, we will use small values of the carrying capacity to determine the role of stochastic fluctuations in the time evolution on 𝕃*. The Gillespie simulations reveal that the initial dynamics of both strands typically follows the deterministic trajectories. Once the dynamics is settled into the quasineutral line, stochastic fluctuations can drive the populations towards one of the corners of the phase space (*x*, *y*), involving the dominance of one of the strands over the other.

Panels (a) and (b) in Fig. 2 display the stochastic dynamics of the strands under two different values of γ and different carrying capacities (*C* = 500 in panel (a); and *C* = 10^3^ in panel (b)). For *C* = 500 and γ = 0.1 (black trajectory in Fig. 2a) the stochastic trajectory follows the deterministic orbit (red dashed line in Fig. 2a) and once it achieves 𝕃*, it evolves towards the corner (0,1) in the phase space, which involves the dominance of the (+) sense strands over the population of (–) strands. A similar result is observed for γ =1 (GR), but the stochastic trajectory ends up in the opposite corner, where (−) strands dominate over the (+) sense strands.

**FIG. 2:**
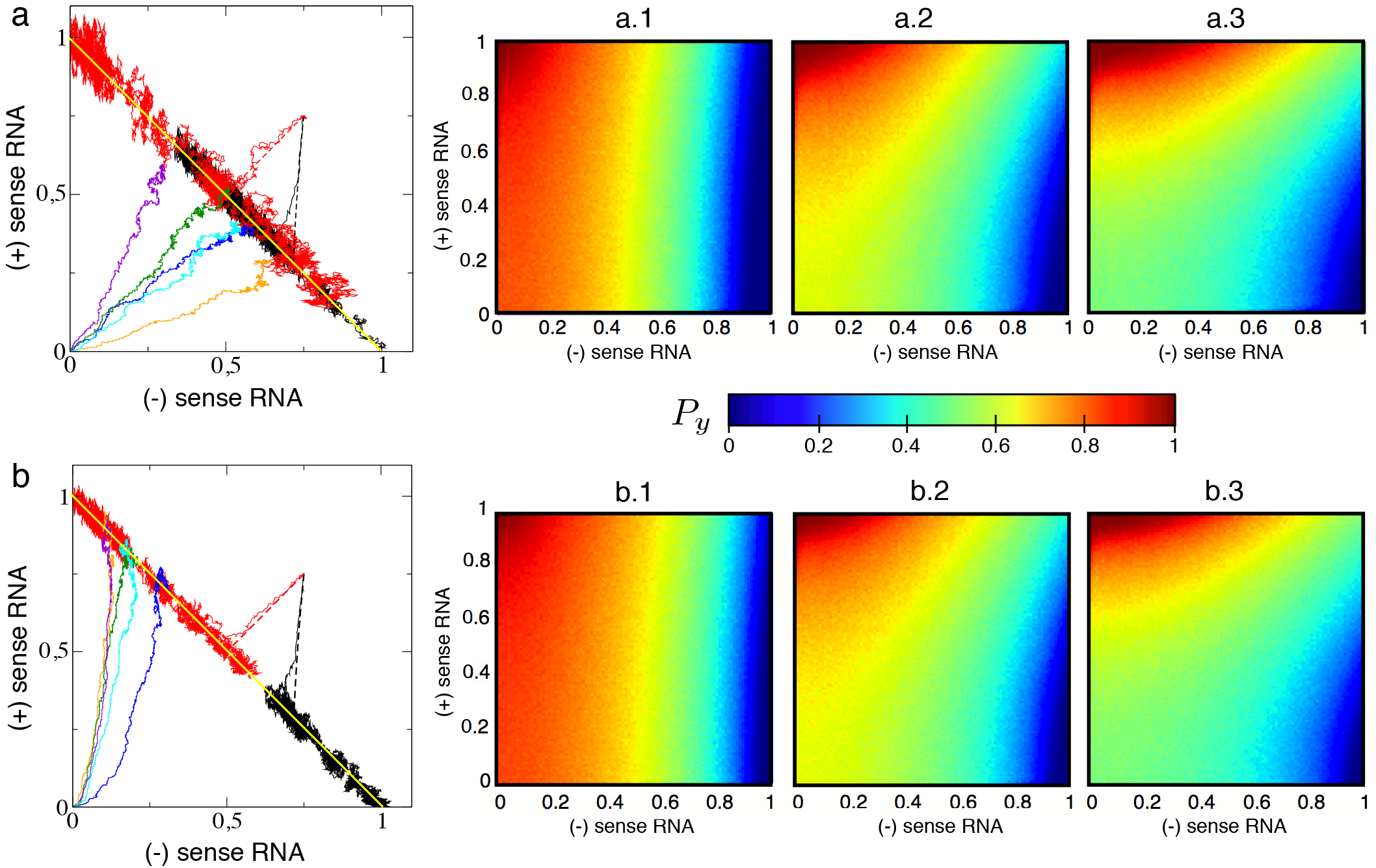
Stochastic dynamics along the quaineutral line 𝕃*. (a) Phase space with stochastic trajectories obtained from the initial condition *m*(0) = *p*(0) = 0.75 using *C* = 500 with γ = 0.1 (black) and γ = 1 (red). The trajectories rapid approach to the line 𝕃* (yellow line) and diffuse along this line due to the stochastic fluctuations. The dashed lines in the phase spaces display the deterministic trajectories obtained numerically also for γ = 0.1 (black) and γ =1 (red) from Eqs. (3)-(4). We also display how several runs from a biologically-meaningful initial condition approach to the line, starting with a single (+) sense and a single (−) sense strand, with γ = 1 (panel (a), with *C* = 500); and γ = 0.5 in (panel (b), with *C* = 10^3^). The small panels at the right display the probability of achieving the state of dominance of positive-sense strands *P*_*y*_ (*z*), with (*x*\* = 0, *y*\* = *1*) using *C* = 500 computed from 5000 replicas using: γ = 0.1 (a.1); γ = 0.5 (a.2); γ = 1 (a.3). (b) Same as in (a) using *C* = 10^3^ and 2500 replicates to compute *P*_*y*_.

The probability of achieving a given state (*x*(*t* ≫ 1) = 1 and *y*(*t* ≫ 1) = 0 or *x*(*t* ≫ 1) = 0 and *y*(*t* ≫ 1) = 1) has been computed as a function of the initial condition within the whole phase space *x* ∈ [0,1] × *y* ∈ [0,1]. Here, for each initial condition sampled regularly in the phase space we have computed the probability *P*_*y*_ from a set of different replicates, computing the number of replicates achieving the state *x*(*t* ≫ 1) = 0 and *y*(*t* ≫ 1) = 1 over the total number of replicates. The results are displayed in panels a.1-3 (using *C* = 500) and panels b.1-3 (setting *C* = 10^3^) in Fig. 2. For replication modes close to the SMR (panels a.1 and b.1) the values of *P*_*y*_ are close to 1 in a wide region of the phase space (orange-red gradient) for *x*(0) ≈ 0.5, displaying the maximum values at the left-upper corner of the phase space, where the initial number of (+) sense strands is large and of (−) sense ones low. As the replication mode approaches to the GR, the region where *P*_*y*_ ≈ 1 remains confined to the corner with initial conditions larger than *y*(0) ≈ 0.6 and *x*(0) ≈ 0.6. This results indicate that under strong stochasticity the SMR model would ensure the production of (+) sense strands, assuming that the initial conditions in the infection of a cell would involve *x*(0) = 0 and *y*(0) ≪ 1.

As mentioned, the dynamics for dynamical systems with a quasineutral line in their deterministic limit involves a fast approach to the line 𝕃*. This feature also happens in the stochastic regime, although once 𝕃* is achieved, noise involves drift along this line. In the previous analyses we have mainly focused on the behavior of trajectories starting outside the invariant line. A way to determine the dynamics on the line is to define a join variable given by *z*_*t*_ = *x*_*t*_ − *y*_*t*_ [14]. By doing so one can simplify the problem and find analytical approximations for the probabilities of achieving one of the possible asymptotic states as a function of *z*. For example, the probability of dominance of (+) sense strands as a function of *z*, labeled *P*_*y*_(*z*). This calculations are developed in the next Section. Before entering into detail on these calculations, we will here estimate *P*_*y*_(*z*) using the Gillespie simulations. The goal of these analyses is twofold. First, we will determine how *P*_*y*_(*z*) behaves for different replication modes. Second, the simulations will allow us to check if the analytical derivations provide a good approximation for *P*_*y*_(*z*) for the system studied in this article.

The probability of achieving the state *z*_*t*_ = −1, with dominance of (+) sense strands, *P*_*y*_(*z*), has been computed for several values of γ, computing the number of replicates achieving *z*_*t*_ = −1 for a given *z* value over a total number of 25 × 10^3^ independent runs. For replication modes close to the SMR, *P*_*y*_(*z*) remains above the antiantidiagonal in the space (*z*, *P*_*y*_(*z*)), meaning that (+) sense strands experience highest fitness (see Fig. 3a with γ = 0.1). This effect is observed also for γ = 0.2, γ = 0.3. For γ ⪆ 0.4 the curve for *P*_*y*_(*z*) starts crossing the antidiagonal, the intersection being at *z* < 0. The increase in γ towards the GR model makes the intersection to happen at *z* = 0. We note that for some values of γ, the curve *P*_*y*_(*z*) also intersects the antidiagonal for values close to *z* = 1 or to *z* = −1. For instance, for γ = 0.1, *P*_*y*_(*z*) intersects the antidiagonal at *z* ≈ 0.7. This phenomenon takes place on the line *z* at the corners of the phase space. In the next Section we discuss and interpret this change in the shape of *P*_*y*_(*z*). The vertical red lines correspond to the theoretical predictions for the crossing values of the antidiagonal by *P*_*y*_(*z*) (see next section for further details).

**FIG. 3:**
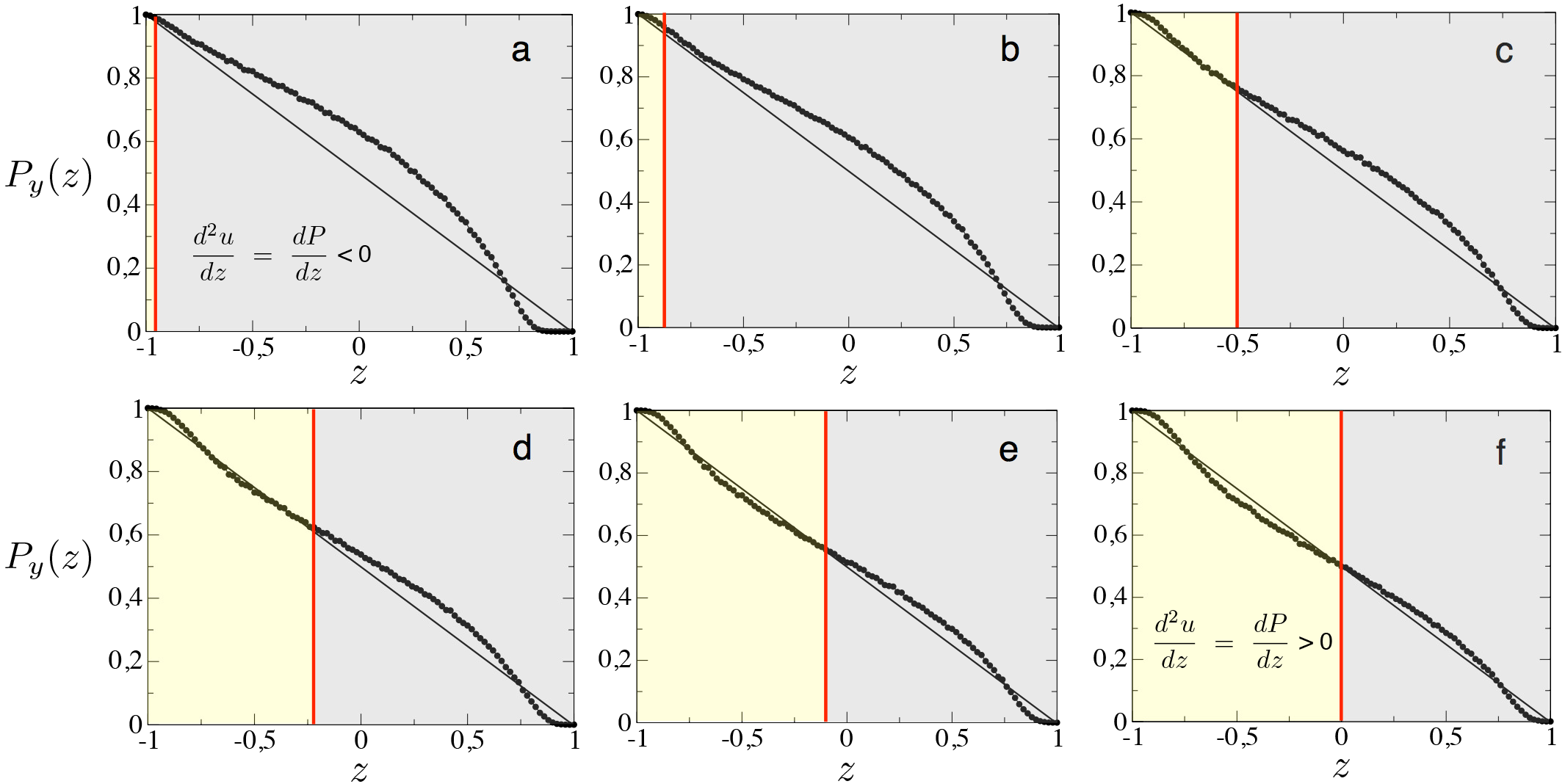
Probability of dominance of (+) sense strands *y*, depending on the value of *z*, given by *P*_*y*_(*z*), obtained from 25 × 10^3^ independent simulation runs using *C* = 500. Panel (a) γ = 0.1; (b) γ = 0.2; (c) γ = 0.4; (d) γ = 0.6; (e) γ = 0.8; and (f) γ = 1. The red vertical lines display the predicted crossing value of *P*_*y*_(*z*) on the antiantidiagonal in the space (*z*, *P*_*y*_(*z*)), given by *z*_0_ in Eq. (24). The *z*_0_ value indicates the change of sign of *d*^2^*u*/*dz* = *dP*/*dz* (see Eq. (23)). For negative values of the second derivative *P*_*y*_(*z*) is convexe (gray regions in panels), while for positive values this function is concave (yellow areas).

Finally, following the approach in [14], we computed the dependence of the mean extinction times (MET, divided by the carrying capacity) on *z* from the Gillespie simulations. These analyses have been performed using two carrying capacities, *C* = 500 and *C* = 10^3^ (Fig. 4). The same results have been obtained for both values of *C*. The METs have been computed from the extinctiontimes (either for *x*_*t*≫1_ = 0 or *y*_*t*≫1_ = 0) obtained from 10^4^ independent runs. Figure 4(a), with *C* = 500 indicates that the METs are symmetric for γ = 1. This means that there is a maximum at *z* = 0 and that for increasing and decreasing *z* values from *z* = 0, the METs decrease similarly (giving place to a parabola-like shape). These times decrease as z grows above or below 0 because the initial conditions approach to the vertices of 𝕃*. As we already saw in some of our analyses, the switch from GR to SMR involves longer extinction times, since the amplification kinetics shifts from purely exponential to subexponential. This can be clearly seen in Fig. 4, where a decrease in γ involves longer METs. Interestingly, the maximum of the parabola displaces towards *z* > 0. This indicates that the METs decrease faster from the maximum of the parabola as *z* → 1 i.e., with larger *x*_0_ values. This phenomenon, arising from the replication model close to the SMR model denotes the fitness advantage of SMR models for (+) sense strands.

**FIG. 4:**
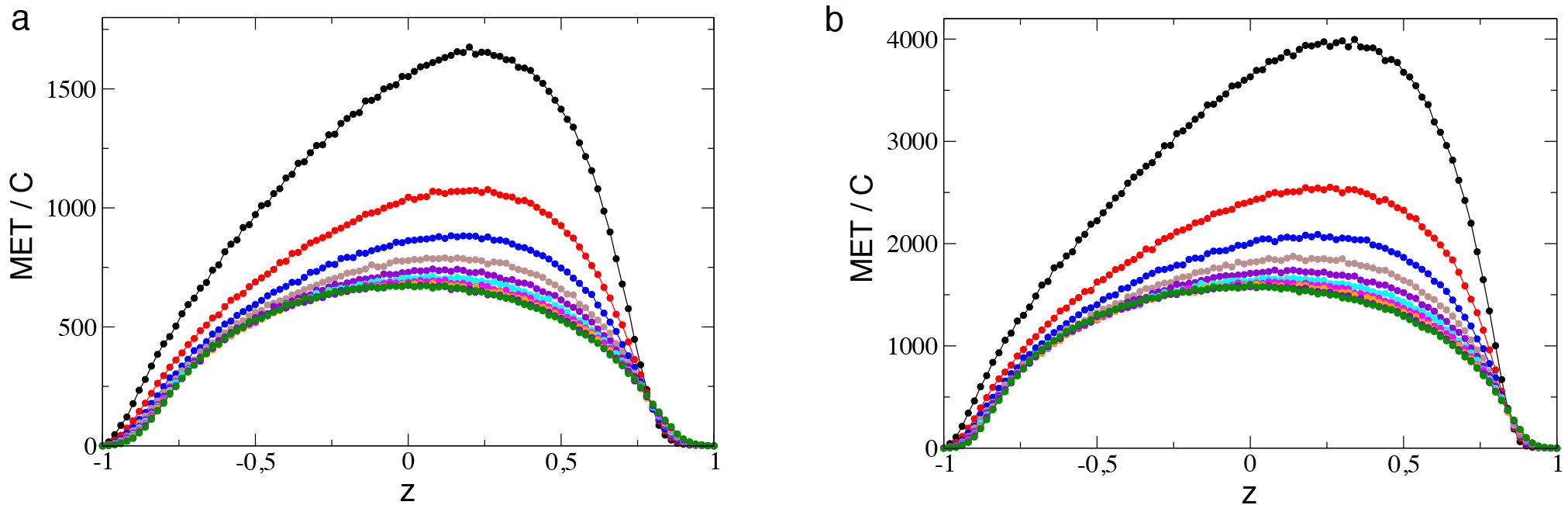
Mean extinction times (MET) divided by the carrying capacity, *C*, as a function of *z* computed from 10^4^ replicas using *C* = 500 (a) and *C* = 10^3^ (b). Ten curves obtained increasing γ are displayed with different colors (from upper to lower): γ = 0.1 (black curve) to γ = 1 (green curve) with γ increments of 0.1.

#### 2. Random dynamics on the quasineutral line

The stochastic dynamics on a quasineutral line of fixed points has been considered previously by both Lin et al. [14] and Kogan et al. [15]. Both approaches are predicated upon the assumption that, once the initial transient during which the system fastly settles on to the quasineutral line, fluctuations away from the quasineutral line quickly relax back to it. By comparison, the dynamics on the quasineutral line, characterized by the variable

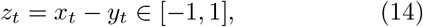

is much slower, and it thus contains the relevant information regarding the long time behaviour of the system. In particular, we here consider the method proposed by Lin et al. [14], who assumed that the relaxation of the fluctuations away from the quasineutral line occurs along the mean-field trajectories. Such assumption allows us to write a reduced dynamics only in terms of the variable *z*_*t*_ (see [14] for a detailed account). The resulting random drift-and-diffusion dynamics proceeds until the system hits one of the two boundaries, *z* = ±1.

Consider an initial condition (*x*_0_, *y*_0_) on the line of fixed points 𝕃*. Under the action of random noise the state of the system is altered so that we can write the new state of the system as (*x*′,*y*′) = (*x*_0_ + *ϕa*, *y*_0_ + *ηb*), where *ϕ* and *η* are random variables such that 〈*ϕ*Ȳ = 〈*η*〉 = 0 and 〈*ϕ*^2^〉 = 〈*η*^2^〉 = 1 [14]. In order to be consistent with the noise terms in the SDEs given by Eq. (13), a and b are so that:

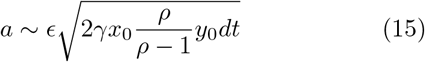

and

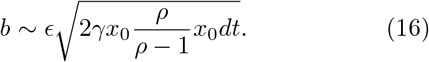

After such perturbation, the system relaxes back to the quasineutral line following the mean-field flow: *x*^2^ − γ*y*^2^ = *cnt*. to a position given by (*x*_0_ − *ξ*, *y*_0_ + *ξ*). Since *z*_*t*_ = *x*_*t*_ − *y*_*t*_, the total displacement along the co-existence line, Δ*z*_*t*_, is given by *x* = −2*ξ*. Therefore, the drift, *υ*(*z*), and diffusion, *D*(*z*), are υ(*z*) = −2〈*ξ*〉/Δ*t* and *D*(*z*) = 4〈*ξ*^2^⟩/Δ*t*, where 〈·〉 represents averaging over the distributions of *η* and *ϕ*. In order to find explicit expressions of 〈*ξ*〉 and 〈*ξ*^2^〉 as a function of the coordinate *z* along the coexistence line, we consider that 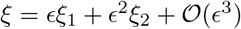 and we use the equation of the mean-field trajectory: 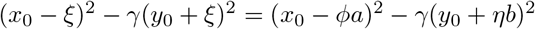. After averaging over the probability distribution functions of *ϕ* and *η*, we obtain that:

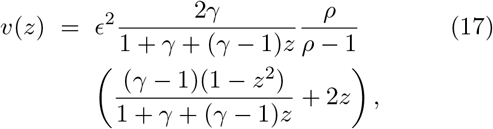

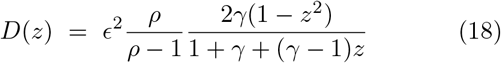

Equations (17) and (18) allow us to write the steady-state backward Kolmogorov equation:

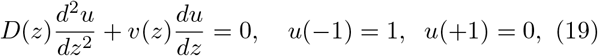

whose solution is the probability of absorption at the boundary *z* = −1 starting from initial condition *z*.

##### a. Measures of fitness

Given the functional form of *υ*(*z*) and *D*(*z*), a full analytical solution of Eq. (19) in closed form becomes cumbersome. It is still possible, however, to define quantities that are informative regarding fitness that can actually be calculated analytically. If the two strands were neutral, the absorption probability would be *u*(*z*) = 1 − *z*. If the actual curve *u*(*z*) is such that *u*(*z*) > 1 − *z*, i.e. the probability of absorption exceeds the neutral value then the fitness of *z* = − 1 is larger than that of *z* =1. If on the contrary, *u*(*z*) < 1 − *z*, the fitness of *z* = −1 is smaller than that of *z* = 1. Since the boundary conditions of Eq. (19) fix the value of *u*(±1), it is straightforward that *u*(*z*) > 1 − *z* if 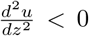 and *u*(*z*) < 1 − *z* if 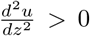. The value of the coordinate *z*_0_ where 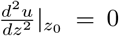 marks the boundary between the regions of higher and lower fitness.

We consider the cases γ = 0 and 0 < γ ≤ 1 separately.

##### b. γ = 0

This case is not biologically meaningful but will help to understand the dynamics of the system under stochasticity. When γ = 0, *x*(*t*) = *cnt* ≡ *x*_0_. The Itô differential equation is then given by

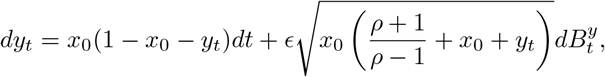

with *y* ∈ [0,1 − *x*_0_]. The drift and diffusion for the reduced process *y*_*t*_ determine the associate statistical features of the competitive exclusion dynamics. Indeed, let 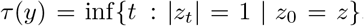. Then the probability that *x*_*t*_ reaches 0 starting from *x*_0_ before *y*_*t*_ starting from *z*, i.e., the probability of domination of the Y-species over the X-species, is

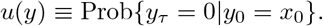

This probability satisfies the boundary value problem

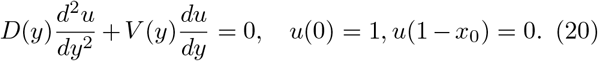

It is easy to see that 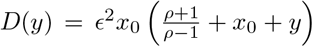 and *V*(*y*) = *x*_0_(1 − *x*_0_ − *y*). Then, we obtain

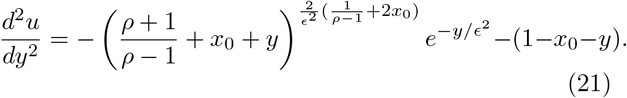

Let us now focus on the case 0 < γ ≤ 1. Here we recover the reduced process obtained with *z*(*t*) in Eq. (14). Since

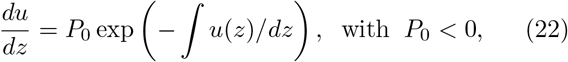

can not be solved analytically, we restrict our calculations to solve *d*^2^*u*/*dz*^2^ = 0. This calculations will allow us to determine whether the curve *P*_*y*_(*z*) is above or below the antiantidiagonal in the space (*z*,*P*_*y*_(*z*)). It can be shown that

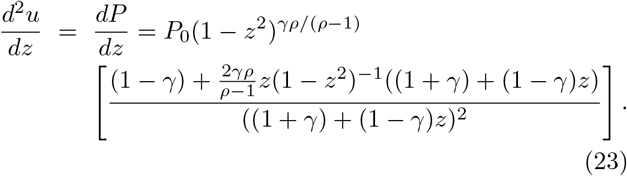

From the previous expression we can obtain the value for which (Eq. 23) crosses the antiantidiagonal in the space (*z*,*P*_*y*_(*z*)) by setting *V*(*z*)/*D*(*z*) = 0. The crossing value *z*_0_ is given by:

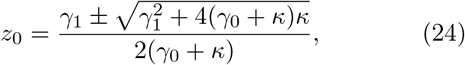

with γ_0_ = γ − 1, γ_1_ = 1 + γ, and 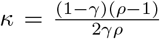 The results for (Eq. 23) are displayed in Fig. 3, overlapped to the outcome of the Gillespie simulations. If we check the prediction provided by *z*_0_ on the crossing point the results display a perfect agreement, since for γ = 1, (Eq. (24)) predicts *P*_*y*_(*z*) crossing at *z* = 0. These predictions hold for decreasing values.

We note that (Eq. 24) predicts two crossing values, although for the range *z* ∈ [1,−1] only one crossing value is found. As mentioned in the previous Sections, the computation of *P*_*y*_(*z*) from the Gillespie simulations resulted in some cases with more crossing values close to the corners (0,1) and (1, 0) of the phase space. Two possible reasons for this deviation from the theory could be given by: (i) a statistical sampling effect in the computation of *P*_*y*_(*z*) in the corners; (ii) a dynamical deviation from the mean-field model.

To check whether hypothesis (i) explains the deviation from the theoretical predictions we have computed again *P*_*y*_(*z*) in the corner 0.5 ≤ *z* ≤ 1 by using 5 × 10^6^ replicates to have more statistical power. These analysis have been carried out for the same values of γ explored in Fig. 3. The same shape of *P*_*y*_(*z*) is found using more replicates. Figure 5a displays the same plot of Fig. 3c (*P*_*y*_(*z*) computed from 25 ×10^3^ replicates for γ = 0.4, and Fig. 5b displays the value of *P*_*y*_(*z*) in the framed region computed from 5 × 10^6^ replicates. Notice that the shape does not change and *P*_*y*_(*z*) crosses the antiantidiagonal. The same results have been obtained using γ = 0.6 (Fig. 5b.1); γ = 0.8 (triangles) and γ1 (open circles) in Fig. 5b.2; and γ = 0.1 and γ = 0.2 (results not shown). The previous results suggest that these deviations are not due to a statistical problem since the crossing of *z* is still found.

**FIG. 5:**
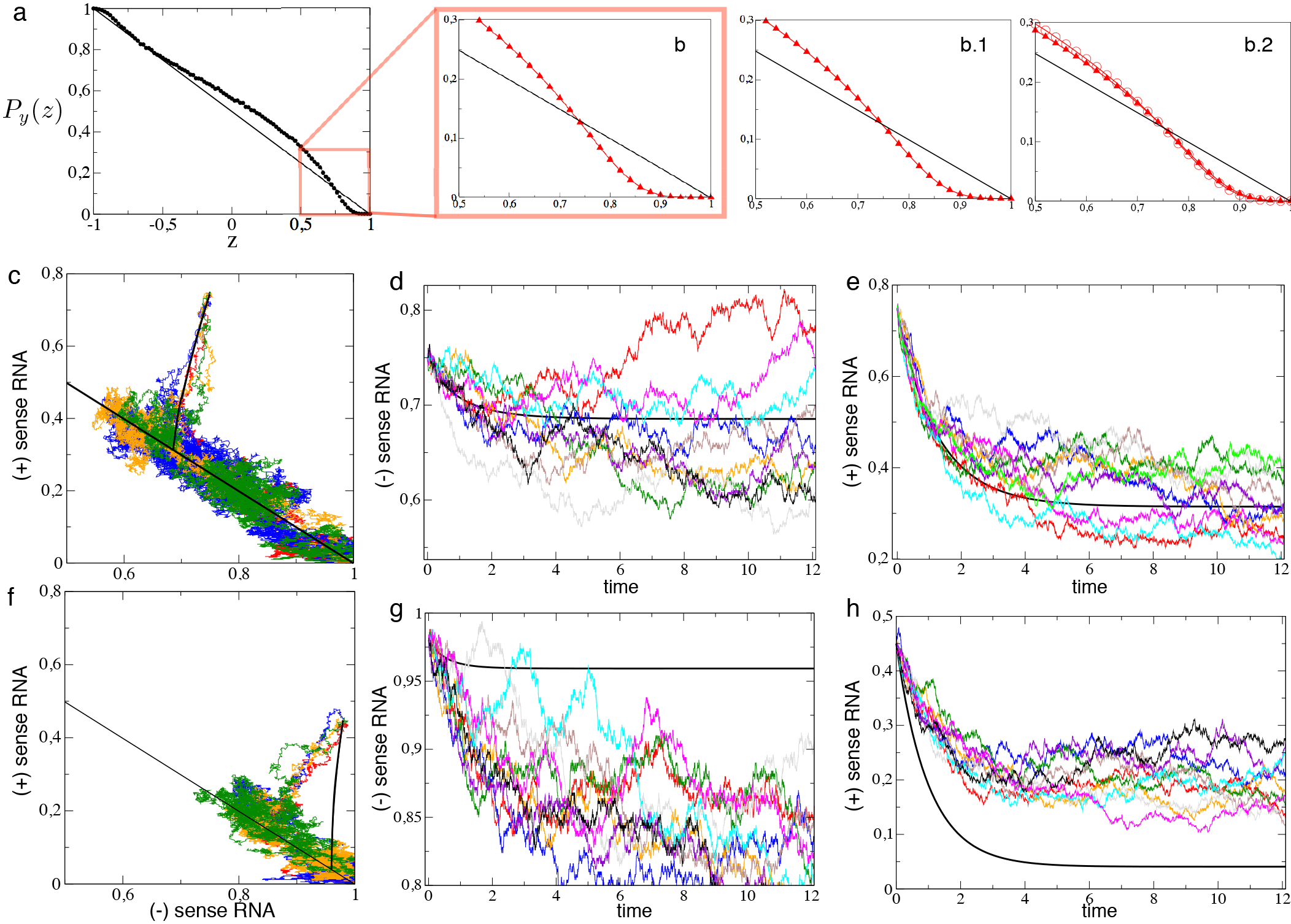
(a) Probability of dominance of (+) sense strands as a function of *z*, *P*_*y*_(*z*), for γ = 0.4 (same panel as in Fig. 3c). Notice that the curve crosses the antiantidiagonal at z ≈ 0.73. (b) Values of *P*_*y*_(*z*) in the region 0.5 ≤ *z* = 1 computed using 5 × 10^6^ replicates. The other two panels show the same results for (b.1) γ = 0.6; and (b.2) γ = 0.8 (triangles) overlapped with γ = 1 (open circles) also using 5 × 10^6^ replicates. Note that increasing the number of replicates to compute *P*_*y*_(*z*) does not involve a change in the shape. (c) Four stochastic trajectories with *x*_0_ = *y*_0_ = 0.75 displayed in the phase space (*x*_*t*_, *y*_*t*_) using γ = 0.2. The solid black line displays the deterministic orbit. In (d) and (e) we display the dynamics of the four trajectories (black, red, dark green, and blue) displayed in (c) plus six other trajectories for (−) and (+) sense strands, respectively. The solid black line also corresponds to the deterministic dynamics. In (f) we display the same as in (c) using *x*_0_ = 0.98 and *y*_0_ = 0.45 as initial conditions also with γ = 0.2. The time series for (−) and (+) sense strands are displayed, respectively, in panels (g) and (h).

According to hypothesis (ii), deviations from the mean-field model trajectories in the corners could explain why the theoretical prediction for *z*_0_ fail. To check this hypothesis we have compared the time dynamics for initial conditions far away and close to the corner of the phase space. The simulations performed using as initial conditions *x*_0_ = *y*_0_ = 0.75 reveal that the stochastic trajectories follow the deterministic dynamics. For example, in Fig. 5(c) we plot four stochastic trajectories in the phase space (*x*_*t*_, *y*_*t*_) overlapped to the deterministic orbit setting γ = 0.2. The stochastic trajectories follow the deterministic dynamics and once they reach the line *z* they fluctuate until the point (0,1) is achieved. The time series displayed in Fig. 5d,e show this good correspondence with the deterministic dynamics. The same results using *x*_0_ = 0.98 and *y*_0_ = 0.45 as initial conditions display a clear deviation from the deterministic dynamics (see Fig.5f and Figs. 5g,h). Note that the stochastic trajectories clearly deviate from the mean-field prediction, suggesting that the theory developed in this Section cannot be applied in models with complementary replicators. This theory was able to explain the quantity *P*_*y*_(*z*) in competing two-species models with self-replication, not with complementary replication, of the form [14]:

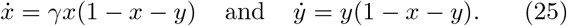

**FIG. 6:**
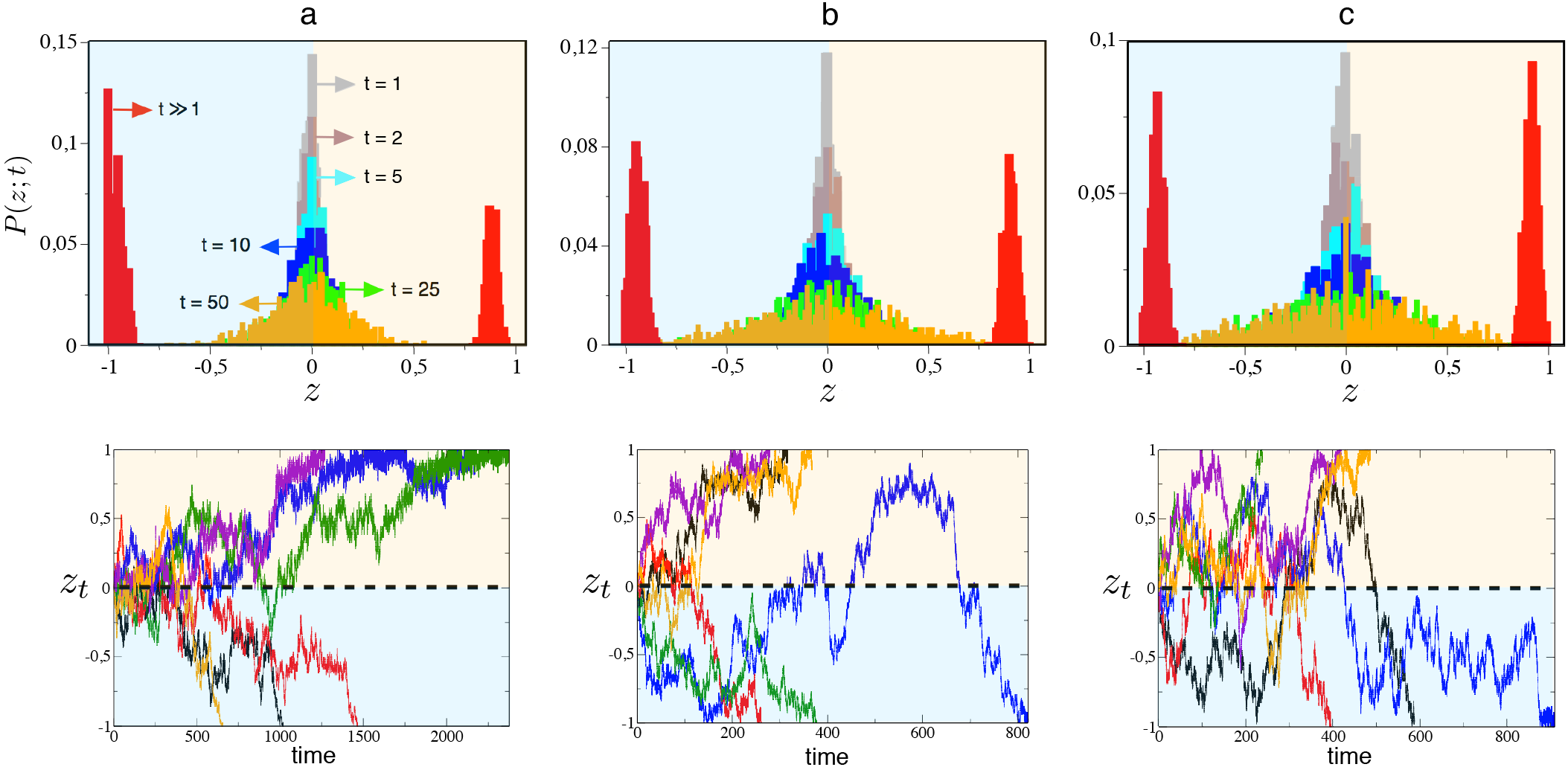
Probability of finding the population of strands at position *z* on the line at time *t*, *P*(*z*; *t*), for different times (indicated with the same colors in the three panels). The histograms display the values of *P*(*z*; *t*) obtained from 10^3^ replicates with initial conditions *x*(0) = *y*(0) = 0.5 for each replicate. Here we set *C* = 500 and: (a) γ = 0.1; (b) γ = 0.5; and (c) γ = 1. Note that along time the replicates split into two possible asymptotic states with full dominance of (+) sense strands (*z*_*t*≫1_ = 1, transparent orange regions) or of (−) sense strands (*z*_*t*≫1_ = 1, blue transparent regions). Below each panel we display six trajectories for *z*_*t*_ starting also at the initial condition *x*(0) = *y*(0) = 0.5.

The theory developed in [14] to compute *P*_*y*_(*z*) was accurate for −1 ≤ *z* ≤ 1 in Eqs. (25).

#### 3. Stochastic dynamics on the quasineutral line drives to noise-induced bistability

Some of our previous analyses indicate the presence of noise-induced bistability in our system. A dynamical view of this phenomena is displayed in Fig. 6, where the fate of multitude of replicates is displayed as a function of time. To perform these analyses we have monitored how different replicates (*n* = 10^3^) evolve in time starting from the same initial condition on *z*_0_ = 0, here setting *x*_0_ = *y*_0_ = 0.5. Then, we have computed for all of the replicates the probability of inhabiting a given region of the line *z* (which has been discretized) at time *t*, probability labeled *P*(*z*; *t*). Figure 6a displays the results for a replication strategy close to the SMR, setting γ = 0.1. Here, for initial times, the population is concentrated at the center of line *z*, and as time advances the distribution of *P*(*z*; *t*) decreases and widens, finally splitting into to well-defined states for which the population can achieve one of the corners of the simplex (red distributions with large *t*). Hence, noise drives the population to two possible asymptotic states.

Below panel a of Fig. 6 we display six trajectories on the line *z*, three of them achieving *z*_*t*_ = 1 and other three achieving *z*_*t*_ = − 1. Note that for γ = 0.1 there is a dominance of *P*(*z*; *t*) towards the region of *z* → −1, indicating that a larger fraction of replicates involve the dominance of (+) sense RNAs, as expected for a replication strategy close to the SMR. This effect is reversed at increasing γ (see Fig. 6b and Fig. 6c which considers GR). In agreement with Fig. 4, the transients for γ = 0.1 are much longer than the ones for the GR model.

## IV. CONCLUSIONS

In this article we have analyzed a simple model considering differences in the mode of RNA replication. Direct [4] and indirect [5–8] evidences from real RNA viruses indicate that they might amplify their genomes following different strategies. On one hand, for single-stranded, (+) sense RNA viruses, the so-called stamping machine replication (SMR) involves that the entire progeny of RNAs is mainly synthesized from one or few (−) sense templates produced after the initial infection with the (+) sense strand. On the other hand, the so-called geometric replication (GR) involves that all synthesized templates have the same chances of becoming templates for further replication. The evolutionary [1, 2] and dynamical [3, 21] implications tied to different replication modes have been mainly investigated using deterministic approaches, although some works have used MonteCarlo simulations to explore the impact of the mode of replication on important evolutionary aspects such as error thresholds [1, 20] or co-infection dynamics [20].

Viral infections are usually initiated by a tiny amount of RNA molecules, and thus demographic fluctuations may have a strong impact on the dynamics of RNA amplification. In this article we have investigated the impact of stochasticity during replication under differential replication modes. Particularly, we have focused on an interesting phenomenon given by so-called quasineutral coexistence. This phenomenon, which involves the presence of a neutral equilibrium state achieved with a fast dynamics, has been described in two-species replicator systems without decay [14] as well as in the competition of strains in disease dynamics [15]. Following the theory developed by Lin and co-workers [14], we provide analytical approximations for the probability of dominance of (+) sense strands. Scenarios favoring the accumulation or the dominance of these strands might result advantageous for (+) sense RNA viruses, which need to package the (+) sense genomes for further infection. Interestingly, we found that such a theory, developed for a model with non-complementary replication, was not accurate enough to describe the stochastic dynamics on the quasineutral line near the boundaries of the phase space in the system studied here.

Our results also reveal a novel type of noise-induced bistability. It is known that random fluctuations can generate novel behavior in dynamical systems, for instance, stochastic resonance [22], noise-induced transitions [23, 24], or the so-called noise-enhanced stabilization [25, 26]. More recent discoveries have provided different mechanisms giving place to the so-called noise-induced bistability. Noise-induced bistability is typically found in dynamical systems which display a single asymptotically stable state in its deterministic limit, but being able to achieve different alternative states due to stochasticity. Different examples of noise-induced stability, which can also be understood in terms of noise-enhanced stability typical from metastable systems (see Refs. [30, 31] for examples on cancer stochastic systems) have been described in many different systems [29, 32–34].

## Acknowledgments

The research leading to these results has received funding from “la Caixa” Foundation. J.S. and T.A. have been partially funded by the CERCA Program of the Generalitat de Catalunya and the MINECO grant MTM2015-71509-C2-1-R.T.A. is also supported by the AGAUR (grant 2014SGR1307) and by a MINECO grant awarded to the Barcelona Graduate School of Mathematics under the “María de Maeztu” Program (grant MDM-2014-0445). S.F.E. has been supported by MINECO-FEDER grant BFU2015-65037-P and by Generalitat Valenciana grant PR0METE0II/2014/021. We want to thank Blai Vidiella for technical help.

